# Targeting *Tmem63b* and *Piezo2* in C-fiber low threshold mechanoreceptor: limitation of *Vglut3*-IRES-*Cre*

**DOI:** 10.1101/2025.07.26.666969

**Authors:** Daniel J. Orlin, Antonio Munoz, Sage Berryman, Destinee Semidey, Swetha E. Murthy

## Abstract

Peripheral somatosensory neurons in the dorsal root ganglia (DRG) transduce mechanical force in the skin and other organs into electrical signals using specialized mechanically activated (MA) ion channels that initiate neuronal activation in response to force. Increasing evidence highlights PIEZO2 as the primary transducer of low-threshold mechanical force in DRG neurons. However, in the absence of *Piezo2*, mice and humans still respond to noxious painful stimuli like pinch, suggesting that additional MA channel(s) likely exist in DRG neurons. Strategies to identify *Cre* lines and DRG subpopulations that select for non-PIEZO2 expressing neurons is therefore an ongoing effort in the field to discover unknown mechanosensors. Here, we investigated a *Vglut3* labeled mouse line as a candidate to identify non-PIEZO2 MA channels in a subtype of DRG neurons called C-fiber low threshold mechanoreceptors (C-LTMRs). Our study carefully demonstrates that the *Vglut3-*IRES*-Cre* mouse line specifically and efficiently labels C-LTMR neurons of the DRG. Electrophysiological recordings using two different *in vitro* mechanical stimulation assays show that the genetically labelled *Vglut3* neurons have robust indentation- and stretch-activated MA currents that are exclusively slowly or ultra-slowly adapting. To determine whether the *Vglut3*-IRES-*Cre* mouse line can be used to delete genes of interest and identify the underlying MA ion channels in C-LTMRs we attempted to generate a *Tmem63b* conditional knockout using this *Cre-*line but detected incomplete loss of *Tmem63b* transcript and lack of TMEM63B- dependent effect on C-LTMR MA currents. Together, our results emphasize that although the *Vglut3-*IRES*-Cre* line is robust in driving expression of a conditional reporter gene, it is inefficient in deleting genes like *Tmem63b* as well as *Piezo2*.

**Statement of Significance:** In this study, we identify a previously uncharacterized *Vglut3*-IRES-*Cre* as a tool to uncover unknown mechanosensors in C-LTMRs of the DRG. We methodically demonstrate that the *Vglut3*-IRES-Cre line labels C-LTMRs and express two mechanosensors, *Tmem63b* and *Piezo2*. However, we were unable to discern which gene contributes to the MA currents due to poor knockout efficiency induced by the *Vglut3-*IRES*-Cre*. Our studies highlight the nuances and challenges associated with *Cre* lines and alert future investigations to exercise caution while using *Cre* lines to discover the contribution of specific genes to mechanosensation.

## Introduction

Dorsal root ganglion (DRG) neurons are primary peripheral sensory neurons that innervate skin and internal organs and detect physical, thermal, and chemical stimuli. DRG neurons are classified into at least 15 distinct subtypes by their transcriptomic profiles and axonal morphology, which underlies their remarkable functional diversity (1–3). One such function is mechanosensation, the process by which neurons are activated by physical forces, giving rise to the sensation of a phone’s vibration or a sibling’s pinch on the arm.

DRG neurons that encode physical stimuli are intrinsically mechanosensitive because they express mechanically activated (MA) ion channels (4). Classically, studies have sorted DRG neurons into discrete categories based on the MA current inactivation kinetics, which range between rapidly (<10ms), intermediately (10-30ms), and slowly and ultra-slowly adapting (>30ms) (5). Through this characterization, a hypothesis developed that each category represents a certain MA ion channel’s activity, which was partially confirmed when research showed that almost all rapidly adapting (RA) currents and some intermediately adapting currents are mediated by PIEZO2 (6,7). Subsequent studies have demonstrated that PIEZO2 is the ion channel that initiates the neuronal response to light touch, gives rise to proprioception, and detects interoceptive cues in both mice and humans (8–12). However, *Piezo2* deficient patients and mice are still able to sense noxious force and deep pressure. Therefore, other MA ion channel(s) likely exist that detect high-threshold mechanical forces and acute mechanical pain.

One strategy to identify the MA channel(s) that responds to high-threshold mechanical force is to isolate DRG neurons with non-RA current, likely not dependent on PIEZO2, and sequence their mRNA to find candidate genes that encode for putative MA channels. Indeed, an in-depth patch-sequencing approach that paired transcriptomics with electrophysiological recordings generated a list of candidates that are enriched in non-RA neurons to be used as a resource in future screens (13). Another approach is to identify DRG subtypes that respond to noxious mechanosensation in the absence of PIEZO2. *In vivo* Ca^2+^ imaging in mouse trigeminal ganglion demonstrated that the DRG neuronal subtype C-fiber low threshold mechanoreceptors (C-LTMRs) respond to gentle and noxious stimuli. However, C-LTMRs retain their responses to noxious stimuli in *Piezo2* conditional knockout (cKO) mice (14). Therefore, C-LTMRs also express a MA ion channel that responds to high-threshold mechanical stimuli, making them sensitive to noxious force.

C-LTMRs have historically been classified in humans by their ability to respond to gentle stroking and to generate a pleasurable sensation (15–17). In addition to encoding affective touch, in mice, studies have also unveiled C-LTMRs’ role in acute mechanical nociception and allodynia (a condition where due to injury or inflammation light touch evokes pain) (18–21). Although it is now known that PIEZO2 is the primary transducer for mechanical allodynia, its contribution to this phenomenon in C-LTMRs is ambiguous (7,22). More recently, the ‘wet dog shake’ phenomenon in hairy mammals was determined to be mediated by C-LTMRs and is dependent on *Piezo2* (23). Most notably the MA currents in C-LTMRs are slowly adapting, which is uncharacteristic of what has been described for PIEZO2 in large diameter DRG neurons and overexpression systems (21,24). Thus, the identity and contribution of the different MA ion channels that underlie C-LTMR properties that could make them sensitive to noxious mechanical forces, remain unknown.

Single cell RNA sequencing database from DRGs indicates that in addition to *Piezo2* the only other known MA ion channel expressed in C-LTMRs is *Tmem63b* (1). TMEM63B, is a monomeric non-selective MA cation channel (25,26). In mice, TMEM63B is responsible for osmotic regulation in outer hair cells, and *TMEM63B* mutations in humans lead to developmental encephalopathies in the central nervous system (27–29). The homolog in *Drosophila*, TMEM63, is crucial for sensory stimuli like humidity sensation and food texture discrimination, and lysosomal function (30–32). While these studies raise the possibility that TMEM63B could function as a sensor for external mechanical stimuli, its role in mammalian mechanosensation remains unknown. Furthermore, the extent of contribution of PIEZO2 and TMEM63B and/or other MA channel(s) to C-LTMR mechanosensation remains to be determined.

Immunostaining approaches have identified unique molecular markers for C-LTMRs like TH, TAFA4, IB4, GNIP, and VGLUT3 (18,21,33,34). Among these, TH and VGLUT3 are commonly used to induce *Cre* recombinase expression in C-LTMRs to genetically label them and to characterize their functional properties. *Th* expression is restricted to C-LTMRs but initiates as early as postnatal day 14 and therefore, *Cre* recombination in this subtype is induced by tamoxifen injections at P14 (1–3,33). However, the TH-2A-*CreER* induces sparse labelling in C-LTMRs and thus has primarily been used to investigate nerve-ending morphology and for *in vivo* calcium imaging (3,33,35). Like *Th, Vglut3* is specific for C-LTMRs after a certain point in development but is expressed broadly during early development (1,2).There are two different mouse lines that drive *Cre* expression under the *Vglut3* promoter, one is a BAC transgenic (*Vlgut3-iCre*) and the other has an IRES2-*Cre* sequence downstream of *Vglut3* (*Vglut3*-IRES-*Cre*) (36,37). The *Vglut3*-*iCre* mouse line labels *Vglut3* lineage cells, some of which exhibit menthol sensitivity and therefore label neurons that express the transient receptor potential cation channel subfamily M member 8 (TRPM8), along with C-LTMRs (18,38). One important distinction to note is that during development *Vglut3* expression turns on in TRPM8 positive neurons, but post development C-LTMRs and TRPM8-expressing nociceptors are distinct populations (23).

In this study we evaluated the utility of the *Vglut3*-IRES-*Cre* line as a C-LTMR marker and its ability to delete MA channels, such as *Tmem63b* and *Piezo2*, to better understand the contribution of *Piezo2* and non-*Piezo2* MA channels in governing C-LTMR mechanosensation. Our results lead to an unexpected discovery that although the *Cre* line labels C-LTMRs, it is inefficient in generating conditional knockouts (cKO) of these two crucial MA ion channels.

## Results

### Validation of *Cre*-mediated recombination in the *Vglut3-*IRES*-Cre* mouse line

To better understand the molecular properties of C-LTMRs and determine the underlying MA ion channels that contribute to their mechanosensation, we sought to identify a mouse line that would label and drive *Cre* expression in C-LTMRs. Although the TH-2A-*CreER* mouse line is considered the most specific for labelling C-LTMRs, we reasoned that the sparse expression of *Cre* and gene deletion only after development in C-LTMRs could present limitations to fully understanding the role of a novel MA ion channel’s effect on development and behavior (33,35). Thus, we chose the *Vglut3-*IRES*-Cre* (*Vglut3^Cre^*), the next most common marker for C-LTMRs (37,39). We crossed *Vglut3^Cre^* mice to the *Cre*-dependent tdTomato reporter line *Ai14* (referred to as *Vglut3^Cre^;Ai14* when pertinent) and determined the specificity and efficiency of the *Cre*-mediated recombination using immunohistochemistry (IHC) by measuring the overlap between tdTomato and DRG neuronal subtypes including medium/large diameter myelinated neurons (neurofilament-200; Nf-200), proprioceptors (Parvalbumin; PV), and C-LTMRs (Tyrosine Hydroxylase; TH) (**Figure 1A-C**). Furthermore, to determine whether, like the *Vglut3*-*iCre* line, the *Vglut3^Cre^* labels TRPM8-expressing nociceptors during development (38,40), we also measured overlap between tdTomato and TRPM8 (**Figure 1F**).

**Figure 1.**
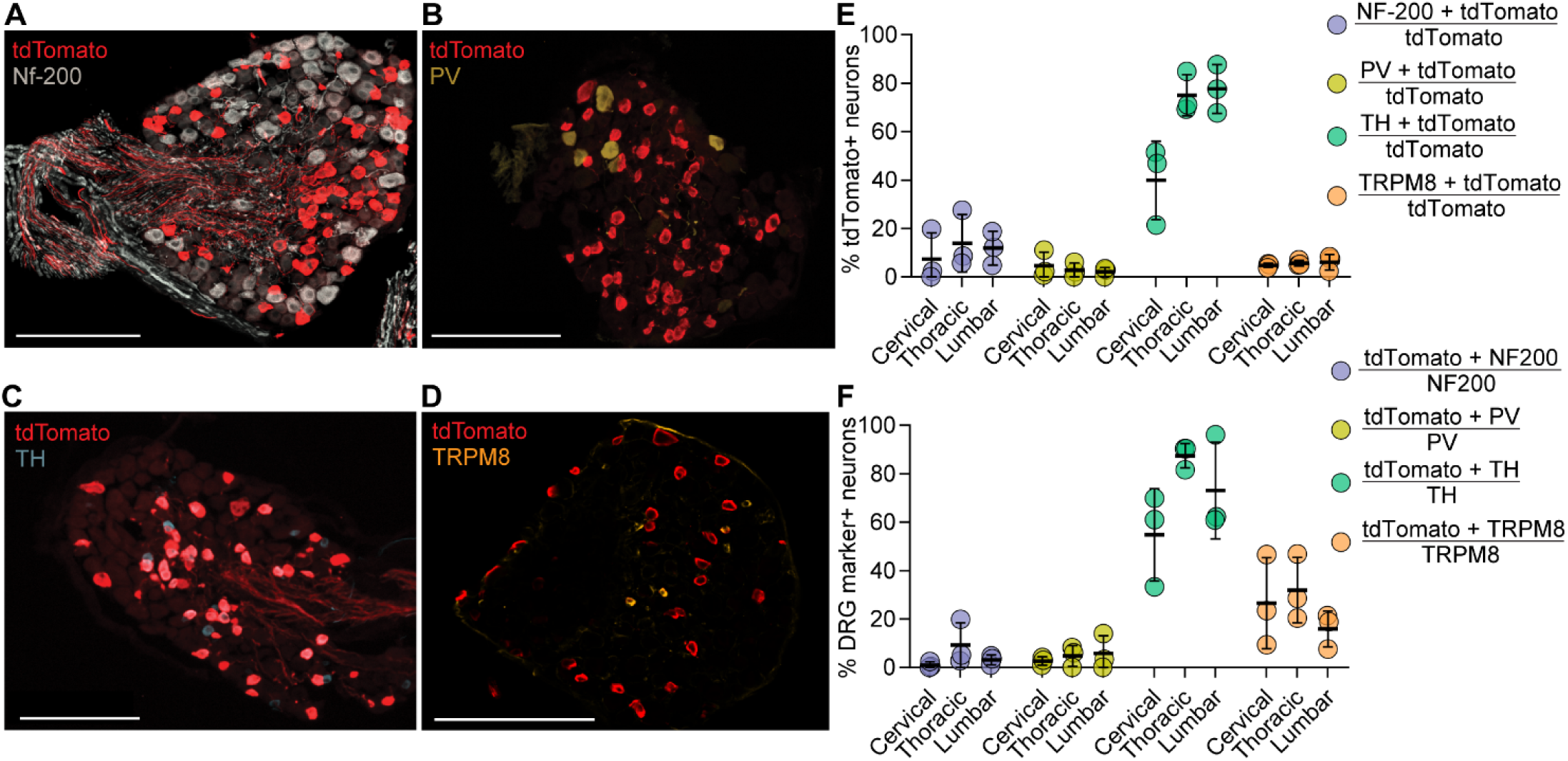
*Vglut3^Cre^*is specific and efficient in genetically labeling C-LTMRs. **A**-**D**: Immunostaining of neuronal cell bodies from *Vglut3^Cre^;Ai14* mice for tdTomato and DRG subtype markers. **A**: Neurofilament-200 (Nf-200), **B**: Parvalbumin (PV), **C:** Tyrosine Hydroxylase (TH), and **D**: TRPM8 (scale bar: 200 µm).**E**: Measure of *Cre* specificity by percentage of tdTomato+ neurons overlapping with each DRG subtype marker divided by the total number of tdTomato+ neurons. **F**: Measure of *Cre* efficiency by percentage of neurons with a DRG subtype marker overlapping with tdTomato, divided by total number of cells for that subtype. Bars represent mean ± S.D.; circles are an average of three images from one animal; N= 3 animals, 3-Females. TRPM8 2- Females, 1-Male.

To measure the specificity of the *Cre*-mediated recombination we calculated the percentage of tdTomato+ neurons that were also labelled by a DRG subtype marker. In lumbar DRGs, we found that 11.9 ± 7.0 % (mean ± SD, N = 3 animals) were myelinated neurons that stained for Nf-200, 2.09 ± 1.84 % were putative proprioceptors, which stained for PV, 77.6 ± 10.0 % of tdTomato+ neurons were C-LTMRs that co-labelled for TH, and 6.07 ± 3.2 % were positive for TRPM8 (**Figure 1E** and **Table 1**). The percentage of overlap between tdTomato+ neurons and DRG subtype markers varied along the rostral-caudal axis with a decrease in TH staining in cervical DRGs (39.9 ± 16.2 %), compared to thoracic (75.0 ± 8.5 %) and lumbar (77.6 ± 10.0 %) DRGs (**Figure 1E** and **Table 1**). To gauge the efficiency of the *Cre*-mediated recombination, we calculated the percentage of neurons with a specific subtype marker that were tdTomato+. In lumbar DRGs, 3.13 ± 2.02 % of Nf-200+ neurons, 5.8 ± 7.27 % of PV+ neurons, and 15.95 ± 7.39 % TRPM8+ neurons were tdTomato+, whereas majority of TH+ neurons were tdTomato+ (73.1 ± 20.0 %); similar values were observed in cervical and thoracic DRGs (**Figure 1F** and **Table 1**). The high degree of overlap between tdTomato and TH suggests that *Vglut3^Cre^;Ai14* efficiently labels C-LTMRs. Additionally, the overlap with TRPM8+ neurons indicate that the *Cre* line labels *Vglut3* lineage cells.

**Table 1.** DRG staining percentages for *Vglut3^Cre^;Ai14* characterization.

| Type of DRG | Nf-200 | TH | PV | TRPM8 |
| --- | --- | --- | --- | --- |
| **Specificity** |  |  |  |  |
| Cervical | 7.38 ± 10.79 | 39.91 ± 16.17 | 4.68 ± 5.63 | 4.78 ± 0.88 |
| Thoracic | 13.91 ± 11.88 | 75.05 ± 8.49 | 2.72 ± 3.08 | 5.47 ± 0.78 |
| Lumbar | 11.93 ± 7.00 | 77.63 ± 10.01 | 2.09 ± 1.84 | 6.07 ± 3.20 |
| **Efficiency** |  |  |  |  |
| Cervical | 0.98 ± 1.32 | 54.76 ± 19.06 | 2.68 ± 1.68 | 26.33 ± 18.25 |
| Thoracic | 9.27 ± 9.22 | 87.45 ± 5.02 | 4.75 ± 4.25 | 31.94 ± 13.50 |
| Lumbar | 3.13 ± 2.02 | 73.08 ± 20.00 | 5.80 ± 7.27 | 15.95 ± 7.39 |
| **Subtype distribution** |  |  |  |  |
| Cervical | 65.07 ± 10.69 | 5.58 ± 2.23 | 14.14 ± 2.85 |  |
| Thoracic | 45.13 ± 9.85 | 21.31 ± 3.00 | 6.41 ± 2.39 |  |
| Lumbar | 45.80 ± 3.89 | 18.67 ± 5.67 | 7.91 ± 3.39 |  |
| **tdTomato** |  |  |  |  |
| Cervical | 10.54 ± 4.95 | 8.10 ± 2.36 | 9.43 ± 3.65 |  |
| Thoracic | 25.48 ± 1.30 | 24.03 ± 1.55 | 24.30 ± 0.86 |  |
| Lumbar | 13.13 ± 3.92 | 14.82 ± 1.99 | 21.25 ± 4.75 |  |

Note: Values indicate mean ± S.D. N = 3 animals and 3 = images.

To rigorously evaluate the *Cre*-mediated recombination in DRG neurons we analyzed the percentage of tdTomato+ neurons across experiments, specifically those used for labeling Nf-200, PV, and TH. We calculated total percentages of labeled neurons with the pan-neuronal label protein gene product 9.5 (PGP9.5) and also evaluated the distribution of both tdTomato+ neurons and the DRG subtypes of interest along the rostral-caudal axis (**Figure S1**). We found the percentage of tdTomato+ neurons remained similar across experiments, suggesting that the *Cre* specificity and efficiency observed, accurately represents *Vglut3* expression (**Figure S1A, B** and **Table 1**). We observed some variability in labelling along the rostral-caudal axis with each experimental group (**Figure S1B**); tdTomato staining was the highest in thoracic DRGs (24.0 ± 1.5 %) compared to cervical (8.10 ± 2.36 %) and lumbar (14.8 ± 2.0 %) regions, suggesting that the percentage of C-LTMRs is higher in this region (**Figure S1B** and **Table 1**). We confirmed this by analyzing the percentage of neurons with subtype markers and observed that TH+ neurons were most abundant in thoracic DRGs (**Figure S1C** and **Table1**). The percentage of TH+ and PV+ neurons that we observed matched previous reports, but we noticed variability in Nf-200 staining along the rostral-caudal axis. Overall, tdTomato staining in the *Vglut3^Cre^;Ai14* line is consistent across staining procedures, experimental trials, and animals, and our reported percentages for each DRG subtype marker agrees with previously published data (33). *Vglut3^Cre^*;*Ai14* specifically labels C-LTMRs and can be used to as a tool to investigate molecular contributions of MA channels to C-LTMR properties.

### Expression of MA ion channels *Tmem63b* and *Piezo2* in C-LTMRs

RNA sequencing studies in mice have shown that C-LTMRs co-express *Tmem63b* and *Piezo2* (*1,2*). We wanted to confirm these observations in our *Vglut3^Cre^;Ai14* mice and further characterize the percentage of C-LTMRs that express both these channels in an attempt to reveal the underlying mechanosensors. To visualize expression, we used *in situ* hybridization (ISH) to label DRG neurons for *Tmem63b* and *Piezo2* transcripts and compared against tdTomato fluorescence (**Figure 2A**). *In situ* images show broad expression for each gene in tdTomato+ and tdTomato-neurons. Within the tdTomato+ cells, 93.7 ± 7.1 % of cells co-express *Tmem63b* and *Piezo2* (**Figure 2B**). We next evaluated TMEM63B protein presence in the DRG. For this we utilized a previously reported *Tmem63b^HA/HA^* mouse that has a haemagglutinin (HA) epitope fused to the N-terminal of TMEM63B in the endogenous locus of the gene, to perform IHC on DRG neurons and compare the expression profile against its localization with the C-LTMR marker, TH (28,33). 100 % of TH- positive neurons were TMEM63B-positive (**Figure 2C**-**D**). These results indicate that TMEM63B is present in C-LTMRs and confirm previously reported RNA sequencing studies, demonstrating that C-LTMRs co-express *Tmem63b* and *Piezo2*.

**Figure 2.**
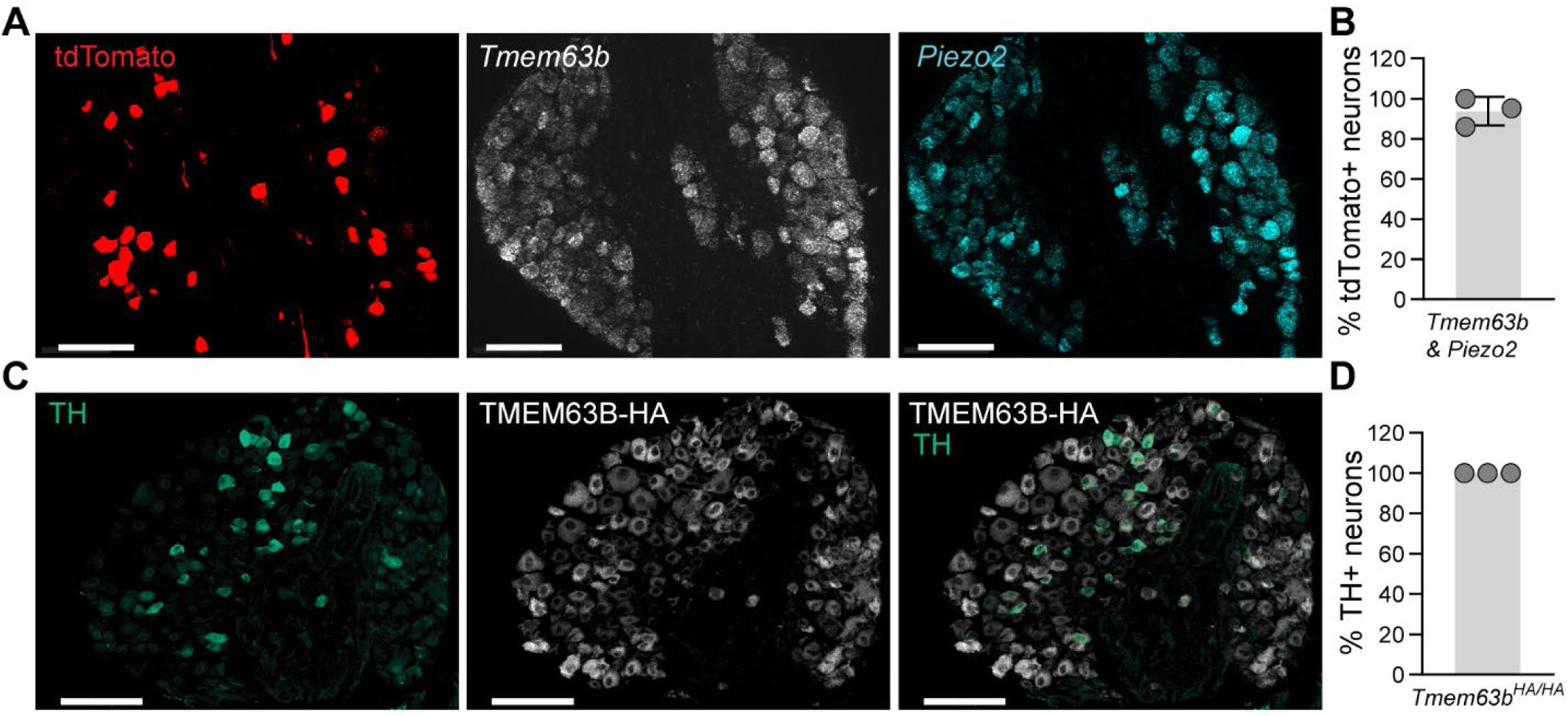
*Tmem63b* and *Piezo2* expression in C-LTMRs. **A**: Thoracic DRG somas from *Vglut3^Cre^;Ai14* mice with *Cre*-dependent tdTomato fluorescence, co-labeled with *in situ* probes against *Tmem63b* and *Piezo2* (scale bar 100 µm). **B**: Percentage of tdTomato+ neurons that co-express both *Tmem63b* and *Piezo2* calculated by mean fluorescence above negative probes (N = 3 animals). **C**: Thoracic DRG somas from *Tmem63b^HA/HA^* mice stained for Tyrosine Hydroxylase (TH), HA-tag, and overlay of both channels (scale bar 100 µm). **D**: Percentage of TH positive DRG neurons with HA staining (N = 3 animals). Bars represent mean ± S.D. Each circle represents an average of three images from one animal.

### Characterization of indentation- and stretch-activated currents in C-LTMRs

We next investigated whether the *Vglut3^Cre^* line can be utilized to generate cKO of *Tmem63b*, to reveal TMEM63B’s contribution to C-LTMR non-*Piezo2* MA currents. We reasoned that if TMEM63B mediates MA currents in C-LTMRs we would observe a change in MA currents of the neurons labelled by *Vglut3^Cre^* and the remaining currents would potentially be mediated by PIEZO2 (14,23). We crossed *Vglut3^Cre^;Ai14* mice to *Tmem63b^fl/fl^* to generate *Tmem63b* cKO animals. For simplicity, we will refer to the wildtype *Vglut3^Cre^;Ai14;Tmem63b^wt/wt^* mice as *Tmem63b^wt^* and the *Tmem63b* cKO *Vglut3^Cre^;Ai14;Tmem63b^fl/fl^* mice as *Tmem63b^fl^*. To first determine whether the *Vglut3^Cre^* labelled neurons recapitulated electrophysiological properties of C-LTMRs, we performed whole-cell current clamp experiments from tdTomato+ neurons in dissociated DRG cultures from *Vglut3^Cre^;Ai14* mice. We were able to distinguish tdTomato+ C-LTMRs from the 13% of tdTomato+ non-C-LTMR neurons based on small soma size and observed action potentials with wide half-width of 2.84 ± 0.81 ms (N = 18), characteristic of C-LTMRs (**Figure 3A**). Further analysis of electrical properties of tdTomato+ neurons from *Tmem63b^wt^* and *Tmem63b^fl^* showed no differences in AP half-width, rheobase, or membrane resistance (**Figure 3A** and **Table 2**).

**Figure 3.**
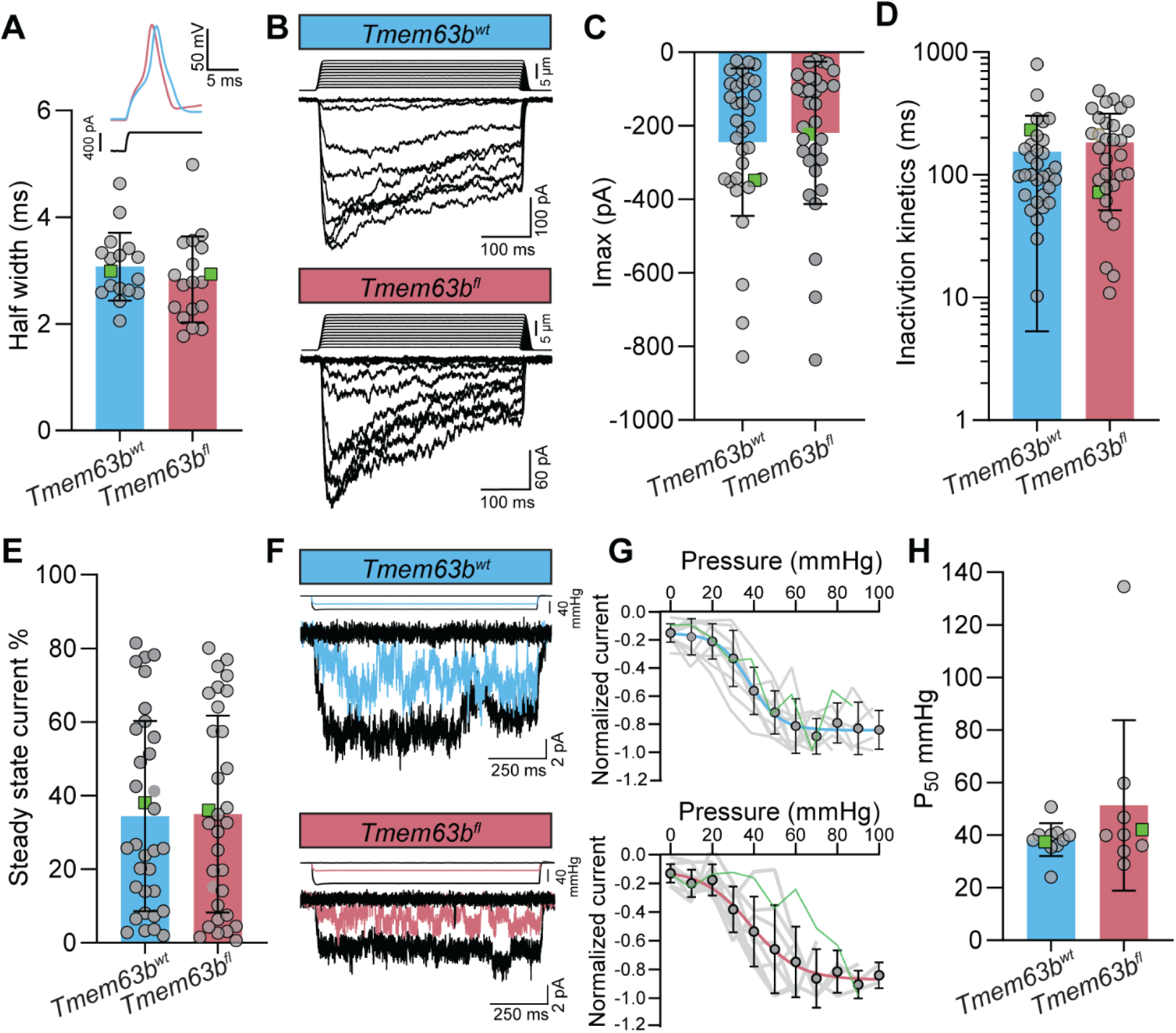
Cultured C-LTMRs have MA currents in response to both indentation and stretch stimuli. **A**: Inset: Representative trace of action potential of a tdTomato+ DRG neuron from *Tmem63b^wt^*(blue) vs *Tmem63b^fl^* (red) mice. Half-width of action potentials within 0-4 pA of rheobase. (*Tmem63b^wt^*; 3.07 ± 0.63 ms, cells = 17 and animals = 4, 2-Males, 2-Females, *Tmem63b^fl^*; 2.84 ±.80 ms, cells= 18 and animals= 5, 3-Males, 2-Females; unpaired ttest *p* = 0.34). **B**: Representative trace of indentation-induced whole-cell MA currents from *Tmem63b^wt^* and Tmem*63^fl^* C-LTMRs. **C**: Maximum peak current (*Tmem63b^wt^*;-244.1 ± 200.4 pA, cells= 33 and animals= 6, 2-Males, 4-Females, *Tmem63b^fl^*; 218.7 ± 193.7 pA, cells = 32 and animals = 6, 1-Male, 5-Females, unpaired t-test *p* = 0.61). **D**: Inactivation kinetics plotted on log_10_ axis (*Tmem63b^wt^*; 154.7 ± 149.4 ms, cells = 32 and animals = 6, 2-Males, 4-Females, *Tmem63b^fl^*; 183.9 ± 132.3 ms, cells = 31 and animals = 6, 1-Male, 5-Females, unpaired t-test *p* = 0.42). **E**: Steady state current taken as a percentage of the peak current (*Tmem63b^wt^*; 34.42 ± 25.82, cells = 33 and animals = 6, 2-Males, 4-Females, *Tmem63b^fl^*; 34.96 ± 26.75, cells = 33 and animals = 6, 1-Male, 5-Females, unpaired t-test *p* = 0.93). **F**: Representative traces of outside-out stretch currents from *Tmem63b^wt^* and *Tmem63b^fl^*. **G**: Current vs pressure relationship, current normalized to maximum response. Shown is a Boltzmann fit of the average values for each group, grey lines are individual cells and green indicates the representative trace’s corresponding measurement (*Tmem63b^wt^*; cells = 11 and animals = 5, *Tmem63b^fl^*; cells = 10 and animals = 4, Boltzmann fit is not different *p* = 0.80). **H**: P_50_ of stretch response fit to a Boltzmann curve (*Tmem63b^wt^*; 38.28 ± 6.24 mmHg, cells = 11 and animals = 3, *Tmem63b^fl^*; 51.32 ± 32.39 mmHg, cells = 9 and animals = 3, unpaired t test *p* = 0.21). **A**, **C**-**E**, **H**: Bars represent mean ± S.D., green squares indicate the example trace’s corresponding measurement.

**Table 2.** Electrical properties of C-LTMRs.

| Electrophysiological Properties | *Tmem63b<sup>wt</sup>* | *Tmem63b<sup>fl</sup>* |
| --- | --- | --- |
| Rheobase (pA) | 181.4 ± 88.8 | 164.2 ± 68.2 |
| Membrane Resistance (Ω cm<sup>2</sup>) | 372.8 ± 151.2 | 381.0 ± 142.6 |
| Half-width (ms) | 3.07 ± 0.63 | 2.83 ± 0.80 |
| Resting membrane potential (mV) | -60.7 ± 9.6 | -56.8 ± 10.6 |

Note: Values indicate mean ± S.D. *Tmem63b<sup>wt</sup>*; 4 animals and 18 cells and *Tmem63b<sup>fl</sup>*; 5 animals and 19 cells.

To activate MA currents in dissociated *Vglut3^Cre^* labelled neurons, we used two distinct methods. In the first method, the soma membrane is indented with a blunt glass probe in whole-cell patch-clamp mode and in the second, a patch of soma membrane is excised and stretched in the outside-out patch-clamp mode. Each of these techniques represents different modes of mechanical stimulation and activation of MA ion channels (41). All recorded cells had a whole-cell indentation response higher than 15 pA (N = 33 cells) (**Figure 3B, C**). We further characterized the indentation MA currents by their inactivation kinetics, measured by fitting an exponential from peak to steady state current of one of the near-saturating traces. Based on the historical categorization of MA currents by their inactivation kinetics, almost all cells had slowly adapting to ultra-slowly adapting MA currents (*5*) (**Figure 3D**). These observations are consistent with previous reports of MA currents from C-LTMRs that were identified using a *TH^2A-CreER^* mouse line (24). Compared to *Tmem63b^wt^* (N = 33 cells), indentation-induced MA currents from *Tmem63b^fl^* (N = 32 cells) dissociated DRG neurons (**Figure 3B**) showed similar mean maximum peak currents (*Tmem63b^wt^*: 244.1 ± 200.4 pA, vs. *Tmem63b^fl^*: 218.7 ± 197.8 pA), inactivation time constants (*Tmem63b^wt^*: 154.7 ± 149.4 ms, vs. *Tmem63b^fl^*: 183.9 ± 132.3 ms), and steady state current (*Tmem63b^wt^*: 34.42 ± 24.82 %, vs. *Tmem63b^fl^*: 34.96 ± 26.75 %) (**Figure 3C**-**E**).

We induced stretch-activated currents in dissociated DRG neurons in the outside-out patch clamp configuration by applying positive pressure to excised patches at 10 mmHg increments (**Figure 3F**). We observed non-inactivating macroscopic currents from tdTomato+ neurons. To obtain half-maximal threshold to activation, P_50_, we normalized stretch-activated currents from individual cells to the maximum response of that patch and fit with a Boltzmann curve (**Figure 3G**). We observed similar P_50_ values of stretch-activated currents from cells of *Tmem63b^wt^* and *Tmem63b^fl^* mice, 38.28 ± 6.24 mmHg (N = 11) and 51.32 ± 32.39 mmHg (N = 9), respectively (**Figure 3G, H**).

Together, the detailed electrophysiological characterization of *Vglut3^Cre^;Ai14* indicates that this mouse line labels a population of DRG neurons, which based on electrical properties confirm are C-LTMRs, are all mechanosensitive, and have a homogenous property of slowly inactivating indentation- and stretch-activated currents. However, no difference in MA current properties were observed between *Tmem63b^wt^* and *Tmem63b^fl^* animals.

### *The Vglut3^Cre^* line does not delete *Tmem63b* or *Piezo2* within C-LTMRs

Expression of *Cre*-mediated tdTomato in the *Vglut3^Cre^*;*Ai14* mouse indicates that the *Cre* is functional; however, the lack of differences in MA currents in the *Tmem63b^fl^* mice compelled us to determine the efficiency of *Tmem63b* knockdown in the cKO mice. To gauge the extent of *Tmem63b* deletion in *Tmem63b^fl^* mouse, we used ISH with a probe that recognizes exon 5 in *Tmem63b*, the region flanked by *loxP* sites in the *Tmem63b^fl^* mice. We performed these experiments in dissociated DRG neurons to recapitulate the conditions used for the electrophysiology experiments. To correlate loss of *Tmem63b* transcript with *Cre* expression, we combined ISH with IHC to stain for *Cre*-dependent tdTomato. Surprisingly, we observed *Tmem63b* puncta in a majority of tdTomato+ neurons in *Tmem63b^wt^* and *Tmem63b^fl^*mice (**Figure 4A, B**). To quantify these observations, we generated 3D regions of interests (ROIs) within all tdTomato cells and counted the *in situ* puncta for each ROI. We binned tdTomato cells by 0-1 puncta for no transcript expression, 2-20 puncta for medium transcript expression, and 21-104 puncta for high transcript expression. 2.8 ± 2.8 % of tdTomato+ neurons in *Tmem63b^wt^* mice had 0-1 *Tmem63b* puncta compared to 12.0 ± 7.0 % of tdTomato+ neurons in *Tmem63b^fl^* mice (**Figure 4C**). 81.3 ± 11.7 % of tdTomato+ neurons in *Tmem63b^wt^* mice had 2-21 *Tmem63b* puncta compared to 68.4 ± 7.0 % of tdTomato+ neurons in *Tmem63b^fl^*mice. Both groups had similar amounts of high expressing tdTomato+ neurons (21-104 puncta). Although we see some tdTomato+ neurons in *Tmem63b^fl^*mice without any *Tmem63b* puncta, a majority of cells have medium to high transcript levels.

**Figure 4.**
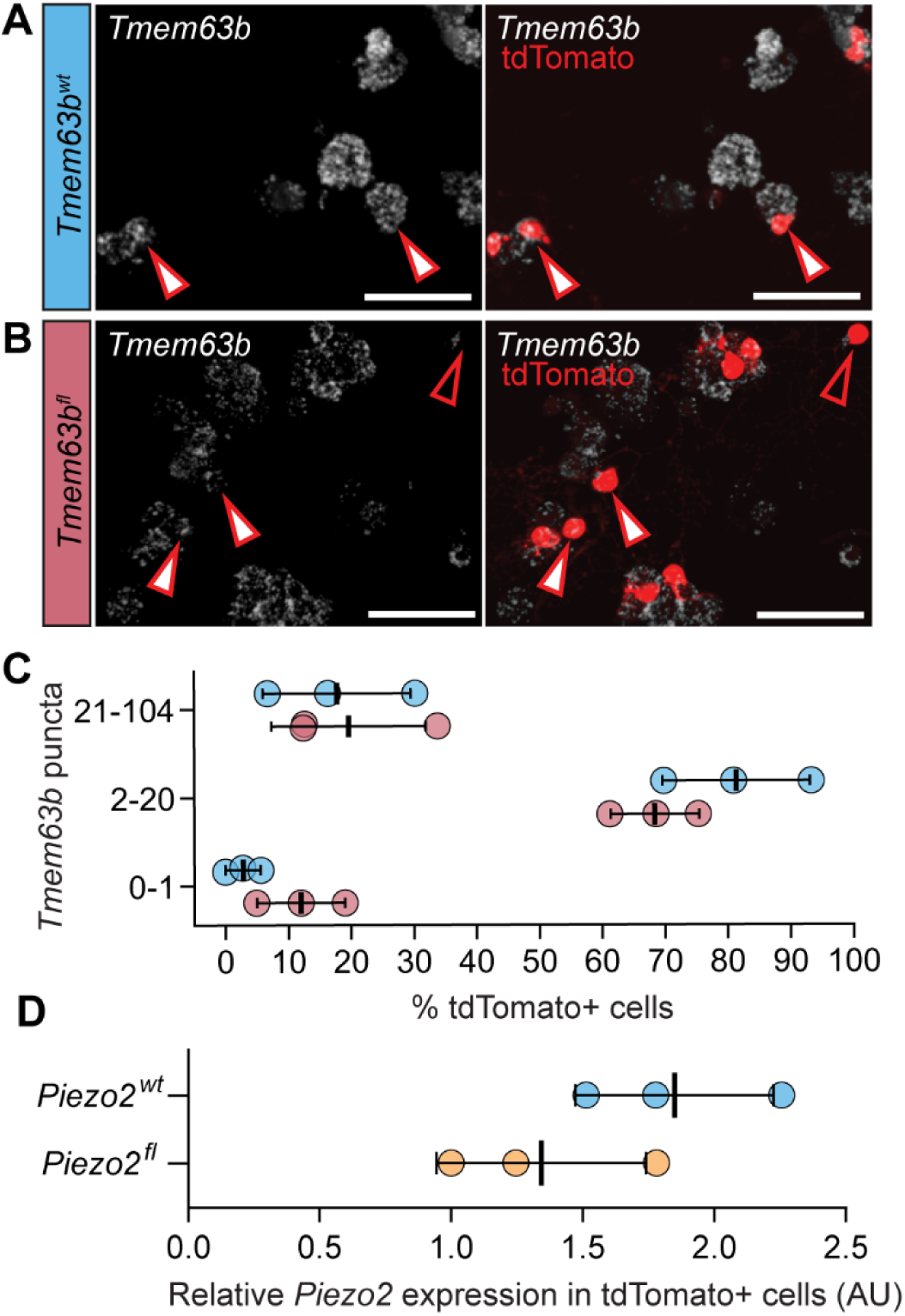
*Tmem63b* and *Piezo2* expression persists in *Vglut3^Cre^* cKO. *In situ* hybridization and immunostaining of cultured DRG neurons from **A:** *Tmem63b^wt^* and **B:** *Tmem63b^fl^,* for probe against *Tmem63b*-exon 5 (white) and tdTomato (red), respectively. Closed arrows denote tdTomato positive neurons with *Tmem63b* puncta. Open arrows denote tdTomato+ neurons without *Tmem63b* puncta. **C**: *Tmem63b expression* measured by the percentage of tdTomato+ cells with a defined number of *in situ* hybridization puncta. Groups binned by the following levels, 0-1 puncta no expression (mean ± SD %), 2-20 puncta medium expression (mean ± SD %), 21-104 puncta high expression (mean ± SD %). *Tmem63b^wt^* (blue), N = 3 animals, 1-Male, 2-Females, vs *Tmem63b^fl^*(red), N = 3 mice, 2-Males, 1-Female. Each dot represents a percentage of the total TdTomato+ cells imaged in a culture from *Tmem63b^wt^* −110, 164, 527 cells and *Tmem63b^fl^* −226, 316, 766. Chi-Square (p < 0.0001 χ^2^ = 22.76, df = 2). **D:** *Piezo2* expression (relative to *Gapdh*) in tdTomato+ DRG neurons from *Piezo2^wt^* (blue, N = 3 animals, 2-Males, 1-Female) and *Piezo2^fl^* (yellow, N = 3 animals, 2-Males, 1-Female) mice (mean ± S.D.). Paired t-test *p* = 0.40.

To further validate that the poor *Cre* efficiency in gene deletion by the *Vglut3^Cre^* line was not unique to *Tmem63b*, we crossed *Vglut3^Cre^*:*Ai14* mice to *Piezo2^fl/fl^* mice and generated *Piezo2* cKO (referred to as *Piezo2^fl^*). To determine *Piezo2* knockdown in the cKO we turned to qPCR, given that the *Piezo2^fl/fl^* line has been previously characterized (8). We identified *Vglut3* specific cells by flow sorting tdTomato+ neurons from dissociated DRGs and compared relative *Piezo2* expression in *Cre*-induced tdTomato labelled neurons from *Piezo2^wt^* and *Piezo2^fl^* mice. Consistent with the *in situ* results for *Tmem63b^fl^* mice, we observed a weak 27.3 ± 38.73 % reduction in *Piezo2* levels in *Piezo2^fl^* mice (*Piezo2^wt^*; 1.8 ± 0.37 and *Piezo2^fl^*; 1.3 ± 0.40, N = 3 animals). Together, these results suggest that although the *Vglut3^Cre^;Ai14* line reliably labelled the C-LTMR positive population by driving recombination of the tdTomato expression gene, the efficiency in deleting *Tmem63b* and *Piezo2* is poor.

## Discussion

In this study, we used the *Vglut3*-IRES-*Cre* line to characterize MA current properties of C-LTMRs (39). While Patil et al., 2018 uses *Vglut3^Cre^* to label C-LTMRs, our study reports the percentage of overlap between other DRG subpopulation markers including TH and for the first time quantifies the specificity and efficiency of the *Cre* line (**Figure 1**). Because we validated that the *Vglut3^Cre^* specifically labelled C-LTMRs, using this *Cre* line to conditionally delete *Tmem63b* or other candidate genes was the most direct approach to test whether the channel was a noxious mechanosensor in these cells. We performed an in-depth characterization of two types of MA currents from dissociated C-LTMRs neurons (**Figure 2**). We observed robust indentation-induced whole-cell MA currents with slowly or ultra-slowly adapting properties. We also report for the first time, to our knowledge, stretch-activated currents from C-LTMRs. Consistent with our results that demonstrate *Piezo2* and *Tmem63b* expression in the *Vglut3*-labelled neuron, the non-inactivating stretch-activated currents resemble PIEZO2 and TMEM63B properties in heterologous expression systems (42). However, despite the robust and consistent nature of *Vglut3^Cre^*line to label C-LTMRs, *Cre* recombination in this line is inefficient and failed to generate cKOs of *Tmem63b* or *Piezo2*.

We note that the *Vglut3^Cre^* used in this study likely labels *Vglut3* lineage cells, because 6.07 ± 3.2 % of the tdTomato+ neurons are also positive for TRPM8 (**Figure 1**). Griffith et al., 2019 showed that the menthol sensitive *Vglut3* lineage neurons have a more depolarized resting membrane potential compared to the menthol insensitive neurons. The tdTomato+ neurons recorded by us have a mean resting membrane potential of −60.7 ± 9.6 mV (**Table 2**), in line with the menthol insensitive *Vglut3* lineage neurons (38). Additionally, TRPM8+ neurons are largely mechanically insensitive (14). Given that almost all tdTomato+ neurons have a MA response, our data most likely excludes TRPM8 positive neurons.

Our results highlight the underlying complexity of using *Cre-Lox* systems to generate cKO animals. Although the *Vglut3^Cre^;Ai14* mouse line genetically labeled C-LTMRs by turning on tdTomato expression, it was inefficient at deleting *Tmem63b* or *Piezo2*. To date, ours is the only study that used the *Vglut3^Cre^* to induce a cKO. Thus, the reason for the inefficient deletion of *Tmem63b* and *Piezo2* is unclear. In some conditional lines, the *loxP* sites are too far apart for efficient recombination, but this is not the case in the *Tmem63b^fl^* animal (43). The *loxP* sites that flank *Tmem63b-*exon 5 are 756 base pairs (bp) apart, which is closer than the *Ai14* reporter strain used in this study, 837 bp (44). The loss of *Tmem63b* transcript in ∼14% of TdTomato positive neurons (**Figure 4**) suggests that the *Tmem63b* floxed mouse line used in our study can be used to induce gene disruption, albeit with low efficiency. Additionally, in Chen et al., 2024, it has been confirmed that removal of *Tmem63b-*exon 5 generates *Tmem63b* null mice and showed that the post birth lethality of *Tmem63b* global KO mice is likely due to its role in lung alveolar epithelial cell function (45). Therefore, the inefficient deletion of *Tmem63b* and *Piezo2* in C-LTMRs may be specific to the *Vglut3^Cre^* line used in this study, or unique to features of the genome in DRG neurons. Future investigations with other *Cre* lines or viral strategies that can selectively delete *Tmem63b* in C-LTMRs will be needed to conclusively determine TMEM63B’s contribution to noxious mechanosensation in C-LTMRs, and how it differs from that of PIEZO2.

A major outstanding question in the somatosensory field is the identity of the MA channel responsible for slowly adapting currents. MA currents from DRG neurons fall into four categories based on their inactivation kinetics, and so far, it is thought that rapidly- and some intermediately adapting currents are PIEZO2 dependent (7,8). Our data demonstrates that C-LTMR neurons are an ideal candidate subtype to screen for the underlying ion channel that elicits slowly adapting currents in DRG neurons. First, all *Vglut3^Cre^* neurons labelled by tdTomato have robust MA currents in response to indentation and second, a majority of the MA currents are slowly adapting. These two properties are uncommon when recording from dissociated DRG neurons. Most subtypes typically have at least 20% non-responders and the MA currents have heterogenous inactivation properties (7). It should be noted that while the MA current properties reported in this study agree with Zhen et al., 2019, our results conflict with Lou et al., 2013, which uses a different *Vglut3-IRES-Cre* mouse line than used in this study, who report MA currents with mixed population of inactivation kinetics as well as non-responders in C-LTMRs (24,46). Regardless, all studies including ours, report slowly adapting currents in the putative C-LTMR population, which has high *Piezo2* expression. Therefore, further studies are needed to assess if PIEZO2 indeed accounts for only rapidly and intermediately adapting MA currents. This notion is also supported by evidence from patch-sequencing data that reveals neurons across the four inactivation groups express *Piezo2* (13). Together, these data imply that PIEZO2 might induce rapidly as well as slowly adapting currents.

PIEZO2 MA currents in C-LTMRs might be modulated by auxiliary proteins like TMC7, resulting in the slowly adapting MA current property. In DRG neurons, *Tmc7* deletion leads to a reduction in non-RA currents, and co-expression of *Tmc7* with *Piezo2* in a heterologous expression system recapitulates the breadth of DRG neuron MA current inactivation properties (47). Similarly, TMEM63B could modulate PIEZO2 MA current properties and the slowly adapting currents in C-LTMRs could be a product of both channels. Conversely, conformational changes in PIEZO2 could influence TMEM63B function, as has been demonstrated for PIEZO1-dependent modulation for K_2_P channels (48,49). In this context it is noteworthy that the *in situ* results in **Figure 3** indicate *Piezo2* and *Tmem63b* are co-expressed in other DRG neuronal populations beyond C-LTMRs. Furthermore, the two channels are co-expressed in non-neuronal cells like bladder endothelial cells (11) and outer hair cells (28,50,51). This co-expression pattern elicits the question of whether the presence of two channels with distinct responses to mechanical stimuli might confer cells with their mechanosensitive properties. Finally, the possibility exists that beyond *Piezo2* and *Tmem63b*, C-LTMRs express other MA ion channels that are yet to be discovered. These PIEZO2-, TMEM63B- dependent and independent possibilities will have to be tested in the future using combinations of *Piezo2* and *Tmem63b* tissue-specific cKO mice.

Our data shows that *Tmem63b* is expressed in many DRG neurons, but its role in mechanosensation beyond C-LTMR function and somatosensory neurons remain largely unknown. In the central nervous system, *Tmem63b* is broadly expressed in different neuronal cell types in certain brain regions like the cerebellum (52). Unexplored *TMEM63B* mutations in humans lead to developmental epileptic encephalopathy, which implies that TMEM63B plays a crucial role in these neurons, but the underlying mechanism is yet to be determined (27). Recent studies are beginning to unravel mechanistic details of the structure and function of TMEM63B and the OSCA/TMEM63 family as MA channels (25,53–56). But as *in vivo* mechanosensors, in outer hair cells TMEM63B activation couples with BK (Ca^2+^-activated K^+^ channels) channels to reduce cell swelling, which could be a ubiquitous mechanism utilized by cell types outside the auditory system (28). These studies and the postnatal lethality of global *Tmem63b* KO mice suggest that TMEM63B could be important for proper cellular function and neuronal development (45). Together they stress the importance of future studies to better understand TMEM63B’s role as a mechanosensor.

## Author Contributions

D.J.O and S.E.M conceived the project. D.J.O performed IHC, *in situ,* microscopy, image processing. D.J.O. and S.B. performed image analysis. S.B. and A.M performed TRPM8 IHC, microscopy, and image processing. D.J.O. and A.M. performed electrophysiology. A.M. performed qPCR. D.J.O., S.B., and D.S. managed genotyping and animal husbandry. D.J.O and S.E.M wrote the manuscript with input from other authors.

## Acknowledgements

We thank Drs. Kevin Wright, Kelly Monk, Eric Schnell, and Mathew Pomaville for insightful feedback on the project. We acknowledge Dr. Stefanie Kaech-Petrie and Felice Kelly for technical support and advice with imaging analysis. We are grateful for the support and resources provided by the OHSU Advanced Light Microscopy core (RRID: SCR_009961) and Flow Cytometry Shared Resource (RRID: SCR_009974). We thank Dr. Julia Halford, Dr. Skyler Jackman, and Amanda Senatore for helpful comments on the manuscript. This work was supported by the T32 training grant (5T32NS007466) to D.J.O and the Silver family foundation grant to S.E.M.

## Materials and Methods

### Animals

All animal procedures were approved by the Institutional Animal Care and Use Committee of Oregon Health and Science University. The *Vglut3-IRES-Cre* mouse was purchased from Jax (028534, RRID:IMSR JAX:028534). The *Ai14* mouse was a generous gift by Dr. Kevin Wright (JAX: **007914**, *B6.Cg-Gt(ROSA)26Sortm14(CAG-tdTomato)Hze/J*. *Piezo2^fl/fl^* mouse was a generous gift from Dr. Ardem Patapoutian. *Tmem63b^HA/HA^* was a generous gift by Dr. Yun Stone Shi and previously published in ref. 33. The *Tmem63b^fl/fl^*mouse was acquired from EMMA (EM:07650) as frozen embryos and rederived at the Scripps Research. First generation mice were crossed to *Flp* (flippase) mice to excise the *LacZ* cassette but to retain the *loxP* sites that flank Exon 5 of *Tmem63b*. Genotyping with PCR detects the first *loxP* site; forward primer: GTTCTTCATATTTCAGGCTTCCTTGCTC, reverse primer: TGCAGTTCCAAAGATGACCAGCAG.

The following crosses were made to generate cKO mice for *Tmem63b* and WT littermate control mice. *Vglut3- IRES-Cre^(Cre/+)^*^;^ *Ai14 Tmem63b^(fl/+)^ X Vglut3-IRES-Cre^(+/+)^*^;^ *Ai14 Tmem63b^(fl/+)^;* cKO mice were *Vglut3-IRES-Cre^(Cre/+)^*^;^ *Ai14 Tmem63b^(fl/fl)^,* and WT mice were *Vglut3-IRES-Cre^(Cre/+)^*^;^ *Ai14 Tmem63b^(+/+)^*.

### Whole DRG dissections

For ISH: Animals were perfused with phosphate buffered saline (PBS) followed by 4% Paraformaldehyde (PFA), and after extracting the spine DRGs were dissected and collected in PBS. After all DRGs were removed, the PBS was replaced with 4% PFA for overnight post-fix. After post-fix DRGs were washed with PBS and placed in 30% sucrose until they sunk or at least 3 days. After cryoprotection in sucrose, DRGs were frozen in blocks of optimal cutting temperature (OCT) compound and sectioned in a Leica CM3050 S cryostat. 12-20 μm sections were placed directly on slides that had been washed with RNAase-away.

For IHC: Dissections were conducted as described above, but DRGs were post-fixed for 30-45 minutes. Following sectioning, sections were placed directly on TOMO slides that were untreated and processed for staining (see below).

### Dissociated DRG cultures

All mice were between the ages of 9-14 weeks old. For electrophysiology and basescope *in situ* hybridization, cervical, thoracic, and lumbar DRGs were extracted and collected in 1ml DMEM/F-12 media. After collection, 1 ml of 12.5mg/ml of collagenase IV (Gibco fisher scientific 17-104-019) pre- dissolved in DMEM/F-12 was added. The DRGs were incubated for 1 hour and then were digested in papain (Worthington) at 10 units/ml for 30 minutes. DRGs were then mechanically dissociated with a fire polished glass Pasteur pipette. The DRG-containing solution was layered above a denser 1.5 mg/ml Bovine serum albumin (BSA) solution, which was made in DMEM/F-12 media with 10% serum. The DRG-BSA layered solution was centrifuged at 80 g for 10 minutes, the supernatant was removed, and the neurons were resuspended in DMEM/F-12 media with 10% serum growth medium supplemented with 100 ng/ml nerve growth factor (NGF), 50 ng/ml Glial cell line-derived neurotrophic factor (GDNF), 50 ng/ml Brain-derived neurotrophic factor (BDNF), 50 ng/ml Neurotrophin-3 (NT- 3), 50 ng/ml Neurotrophin-4 (NT-4) and plated on coverslips for electrophysiology experiments or on Lab-Tek II chamber slide that have been treated with laminin.

### *In situ* hybridization

*In situ* hybridization on DRG cryosection was done following the RNAscope multiplex fluorescent assay kit user manual methods from ACDbio, using the 431531 probe for *Tmem63b* and the 400191-c2 probe for *Piezo2*. For each animal the same tissue was used on different slides to run the 3-plex negative control probes and matched by the opal channels used for our experimental probes.

For *in situ* analysis of cultured DRGs, the BaseScope Detection Reagent Kit v2-Red user manual was used. Cultured cells were stained with either the positive, negative, or 1039041-C1 *Tmem63b* BaseScope probe. Immediately after BaseScope staining was completed slides were treated with blocking solution with PBS, 0.1% triton, and 10% donkey serum for 1 hour. Then IHC was performed with the Goat-tdTomato antibody and corresponding secondary antibody.

### Immunohistochemistry

Using a pap pen a barrier was drawn around all DRG sections. Slides were incubated for 1 hour in permeabilization and blocking buffer (PBS, 0.2% triton, and 5% donkey serum). Primary antibodies were diluted in the same permeabilization and blocking buffer and added to incubate overnight at 4°C. The next day primary antibodies were removed, and slides were washed 3 times in PBS for 10 minutes or longer. Secondary antibodies diluted in permeabilization and blocking buffer and were added for 45 minutes at room temperature. After the secondary antibody solution was removed, the slides were washed 5 times with PBS, with the 2^nd^ wash containing 1:5000 DAPI. Slides rested at room temperature until they were dry (1-2 hours), and then coverslips were mounted with Fluoromount-G with DAPI (Fisher 50-112-8966).

### Primary antibodies

**HA/TH staining:** Rabbit anti-HA Cell Signaling Technology #3724 1:250, sheep anti-tyrosine hydroxylase Sigma AB1542 1:1000.

**PGP/tdTomato/DRGmarker**: Guninea pig anti PGP9.5 from Neuromics inc GP 1410450UL at 1:400, goat anti tdTomato OriGene Technologies AB8181-200 1:500, rabbit anti PV GeneTex GTX132759 1:150, rabbit anti TH 1:500 Sigma AB 152, rabbit anti NF-200 1:1000 EMD Milipore ABN76, and rabbit anti TRPM8 1:500 Origene TA423282.

### Secondaries antibodies

**HA/TH staining:** anti-rabbit IgG-Alexa Fluor 555 Biotium 20038 1:500, and anti-sheep Alexa Fluor 647 Jacksom ImmunoResearch 713-605-003 1:500.

**PGP/tdTomato/DRGmarker**: Anti-Guinea pig-cy3 Jackson ImmunoResearch 706-165-148 1:500, Anti-Goat IgG-Alexa Fluor 647 Jackson ImmunoResearch 705-605-003 1:500, and anti-Rabbit IgG-Alexa Fluor 488 Thermo Scientific R37118 or A-21206 1:1000, 2 drops per 1mL of permeabilization and blocking buffer (as per the manufacturer’s instructions).

### Imaging and analysis

***Tmem63b* and tdTomato ISH and IHC Imaging:** Images were taken on a BC43 Andor Spinning Disk Confocal using a 20X objective to obtain a montage of images and z-stacks of almost the entire culture of DRG neurons. **Processing:** The montage of images was loaded into Imaris stitching software to make a single stitched imaged. That image was converted into an arivis file. An arivis pipeline was developed to process and analyze the image. Steps include denoising for all channels, and a background correction for the basescope channel with method: morphology, filter: Preserve bright objects: and radius 1.24 um. **Segmentation:** Part of the arivis pipeline uses a ‘machine learning segmenter’ that was trained with previous images to generate 3- dimentional Regions of Interests (ROI) within tdTomato neurons. Then a blob finder is used to capture ROIs for the Basescope puncta. **Thresholding:** All blobs with 35% of their area in a tdTomato+ cell were counted as puncta. Before the final tally all blobs in the negative control stain within tdTomato neurons were measured for their mean fluorescence and a threshold was made using the 99^th^ percentile of these negative blobs. That threshold was applied to the experimental blobs and any blob with a mean fluorescence below that threshold was discounted. Then each tdTomato+ cell had their blobs recorded and binned by the number of blobs.

***Tmem63b* and *Piezo2* ISH: Imaging:** Following RNAscope, Z-stack images were taken on a ApoTome2 Zeiss Microscope, tdTomato was imaged using its natural fluorescence. **Processing**: Images were processed in Zen Lite by clicking on the ApoTome tab and creating a new image using optical sectioning. The stack was then opened in Fiji where a central z-slice was selected for analysis. **Segmentation:** ROIs were created by using the wand tool (mode set to legacy, tolerance set to 331), and clicking on tdTomato+ cells. Within each ROI the mean intensity of pixels in the ISH channels was recorded. This analysis was the same for all RNAscope images including the negative controls. **Thresholding:** The mean fluorescence from the negative controls was used to create a threshold to determine if a cell expresses *Tmem63b* or *Piezo2*. For each animal all the tdTomato+ cells’ mean fluorescence in the negative control c1 and c2 channels was recorded to calculate the 95^th^ percentile. The 95^th^ percentile calculated for each animal was used as threshold for the corresponding channel from that animals i.e. if the 95 percentile for the c1 channel in animal 1 was 22 any cells from that corresponding channel, *Tmem63b,* in the experimental group that falls below 22 were determined not to express *Tmem63b* (Figure S1A-B).**HA and TH IHC: Imaging:** Images were taken on a BC43 Andor Spinning Disk Confocal with a 20X objective with z-stacks encompassing the entire slice. Each animal had 3 image stacks recorded. **Segmentation:** Analysis was done in Imaris where surfaces were created in the TH channel that represented each TH positive cell. **Thresholding**: The mean fluorescence of TMEM63B-HA within each surface was exported to excel. In prism the 99^th^ percentile was calculated using all the values for the mean fluorescence of the TMEM63B-HA channel within TH positives cells from a *Tmem63b^WT/WT^* animal. That 99^th^ percentile was used as a threshold for TMEM63B-HA staining. If a TH cell had a mean fluorescence above that threshold it was counted as TMEM63B-HA positive cell.

***Vglut3^cre^;Ai14* IHC: Imaging:** Images were taken on a ApoTome2 Zeiss Microscope. **Processing**: Images were processed in Zen Lite by clicking on the ApoTome tab and creating a new image using optical sectioning. Images were then imported into Imaris. **Segmentation:** Using the surface function and some customization, a unique surface program was created for each stain (i.e., the TH stain and NF stain had different parameters). Parameters remained constant within an animal but were adjusted slightly between animals. The PGP 9.5 surfaces were generated first for every image, and then those surfaces were deleted by hand if they did not correspond to a cell. Then the surfaces for the other markers were generated and any surface that did not overlay with the PGP 9.5 surfaces was deleted by hand. **Thresholding**: Using the coloc function a new channel was created that consisted of the overlap between tdTomato and the DRG subtype marker used, with the intensity for each channel staying consistent within animals. A surface was created for that channel as the overlap between the two. Surfaces were then tallied and recorded for analysis in GraphPad (Figure S1C).

**TRPM8 and tdTomato ISH and IHC Imaging:** Images were taken on a BC43 Andor Spinning Disk Confocal with a 20X objective with z-stacks encompassing the entire slice and analysis was done in Imaris.

**Segmentation:** Using the surfaces function in Imaris, a pipeline was developed to label tdTomato+ and TRPM8+ DRG cells and was specific to each animal. Surfaces representing TRPM8+ and tdTomato+ cells were generated initially based on mean channel intensity and then filtered for object area and volume, to exclude nerve fibers and debris. **Thresholding:** The coloc function was used to measure overlap of average TRPM8 intensity in tdTomato+ labeled cells. Thresholds were set on a per-animal basis. Counts of TRPM8+/tdTomato-, TRPM8-/tdTomato+, and TRPM8+/tdTomato+ cells were recorded for each image and summed for each animal (Table 1).

### RT-qPCR to measure *Piezo2* knockdown in *Vglut3^Cre^;Ai14* mice

DRGs were harvested from either *Vglut3^Cre^;*Ai14;*Piezo2^wt/wt^*or *Vglut3^Cre^;*Ai14;*Piezo2^fl/fl^* mice and cultured as mentioned above until the resuspension step. Dissociated neurons were transferred to DPBS buffer containing 2% FBS and 0.01 mM EDTA. The neurons were flow sorted (OHSU Flow Cytometry Core) to isolate tdTomato+ neurons. Sorted neurons were collected in 1X RNA wash buffer and RNA was isolated using the Zymo RNA miniprep kit (catalog No. R1057). cDNA was synthesized from isolated RNA using the High-Capacity cDNA Reverse Transcription Kit (Thermo 4368814) with equal ng amounts of starting RNA. Synthesized cDNA was then used to perform qPCR using predesigned Taqman gene expression assays (Thermo: *mmGapdh*-VIC, 4331182_Mm99999915_g1; *mmPiezo2*-FAM, 4448892_Mm01262434_m1) using a CFX Connect Real-time system (Bio-rad). *Piezo2* transcript levels were analyzed alongside *Gapdh* transcript levels as a loading control, and results for gene expression were computed using the built-in software from Bio-rad.

### Electrophysiology

Extracellular Solution: NaCl 133 mM, KCl 3 mM, CaCl_2_ 2.5 mM, MgCl_2_ 1 mM, HEPES 10 mM, Glucose 10 mM, pH adjusted to 7.3 with NaOH, then osmolarity adjusted, if necessary, with Mannitol solution to 310 mOsm.

Intracellular solution: KCl 133 mM, EGTA 5 mM, CaCl_2_ 1 mM, MgCl_2_ 1 mM, HEPES 10 mM, pH adjusted to 7.3 with KOH, then osmolarity adjusted, if necessary, with Mannitol solution to 300 mOsm. Prior to use, Mg-ATP is added at 4mM and Na-GTP is added at 0.4mM. Patch pipettes were made from borosilicate glass O.D.:1.5mm I.D.:0.86mm pulled on a Sutter P-97 puller and polished with a microforge MF-830.

To identify C-LTMRs in dissociated DRG neuronal cultures we used tdTomato and small cell soma size. Soma size allowed us to distinguish C-LTMRs from the ∼13% of larger diameter non-CLTMR neurons labelled by tdTomato in the *Vglut3^Cre^;Ai14* line (18). To confirm that we could reliably identify C-LTMRs based on their small soma size, we turned to the inherent electrical properties of C-LTMRs defined by their wide action potential. Most DRG neurons have action potentials with sub-millisecond half-widths, whereas C-LTMRs have action potentials with a half-width wider than 1.5 ms (24). Indeed, the mean half-width of an action potential recorded from the tdTomato+ neurons was 3.07 ± 0.63 ms (N= 17) as reported in **Figure 3**.

Whole-cell patch clamp: Currents were recorded in whole-cell voltage clamp mode using Axopatch 200b amplifier (Molecular devices) sampled at 10 Hz and filtered at 2 kHz. Pipettes used were between 1.7-4.5 MΩ. Cells for patching were chosen based on their tdTomato fluorescence and small soma size. Cells were held at −60mV and were stimulated by a blunt glass probe positioned at an angle ∼80° with a piezo-electric crystal microstage (E625 LVPZT Controller/Amplifier; Physik Instrumente). The probe was typically positioned ∼1-2 μm from the cell body, stimulus duration was 400ms and steps were applied at an increment of 1 μm per sweep.

Outside-out patch clamp: Set up for outside-out recording was same as whole-cell patch clamp recordings Stretch-activated currents in the outside-out patch was induced by applying positive pressure to the excised membrane patch with a High-Speed Pressure Clamp 2-SB by ALA science. Pressure stimulus duration was for 1 sec, and steps were applied at increments of 10 mmHg per sweep. Cells were deemed as non-responder if the max current response did not exceed the mean max current response of all cells at 0 mmHg because this was equal to the noise of the baseline current.

Current clamp experiments: Currents were recorded in whole-cell current-clamp mode using Axopatch 200b amplifier (Molecular devices) sampled at 10 Hz and filtered at 2 kHz. Neurons were recorded in the extracellular bath solution. Pipettes used were between 3-6 MΩ. After a giga-ohm seal was formed, pipette capacitance was offset. Current was injected into the soma to maintain a membrane potential of 60 +/− 5 mV. In rheobase recordings, a 20 pA step stimulus was applied for 200 ms, every 10 seconds until a cell fired an action potential. Membrane resistance was calculated using Ohm’s law with data from the rheobase recordings. The amount of injected current, average resting membrane potential prior to the current stimulus, and average change in membrane voltage during current stimulus were recorded and these values were used to calculate membrane resistance. Half width was measured from action potentials elicited at a saturating stimulus of 400 pA for 200 ms. Cells were excluded from analysis if, after forming a giga-ohm seal the break-in resting membrane potential of a cell was above −40 mV, or if during rheobase recordings the cells failed to elicit an action potential past 0 mV after applying more than 400 pA of current, or if more than 100 pA of current was injected during recordings to maintain membrane potential of 60 +/− 5 mV.

All electrophysiology data was analyzed in Clampfit11.1 and GraphPad Prism.

---

**Supplementary Figure 1.**
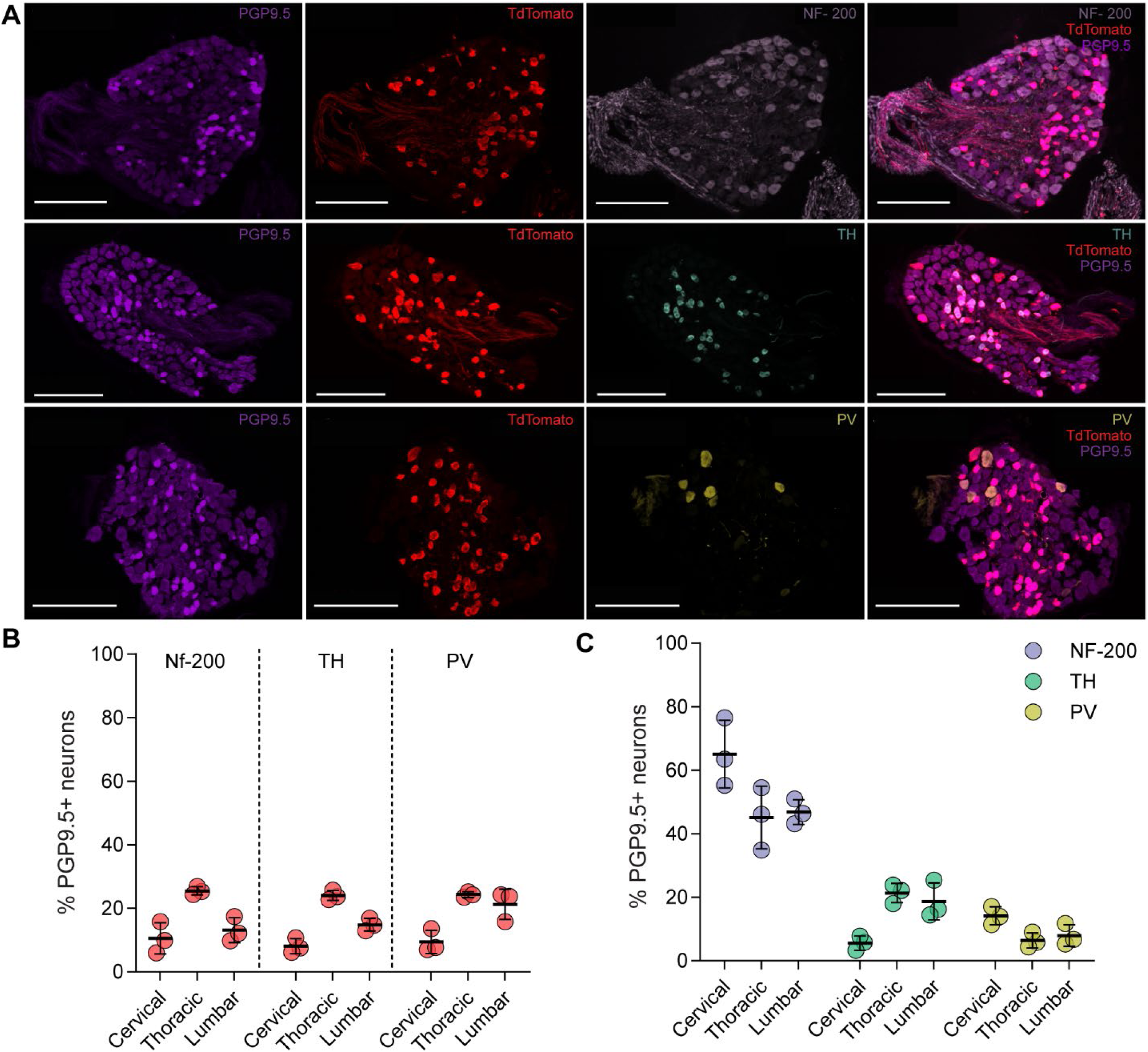
Staining of subtype markers and tdTomato is consistent across animals and experiments. **A**: Immunostaining of neuronal cell bodies from *Vglut3^Cre^;Ai14* mice stained for PGP9.5, tdTomato, and NF-200 (top), TH (middle), or PV (bottom); overlap of all channels are also indicated (scale bar: 200µm). **B**: Percent of tdTomato neurons by calculating the overlap of tdTomato and PGP9.5 positive neurons and dividing it by the total neurons (PGP9.5 positive), separated by DRG subtype marker staining. **C**: Quantification of subtype abundance as a percent of PGP9.5 positive neurons that overlap with a DRG subtype marker. Bars represent mean ± S.D. ; circles are an average of three images from one animal; N=3 Animals, 3-Females.

**Supplementary Figure 2.**
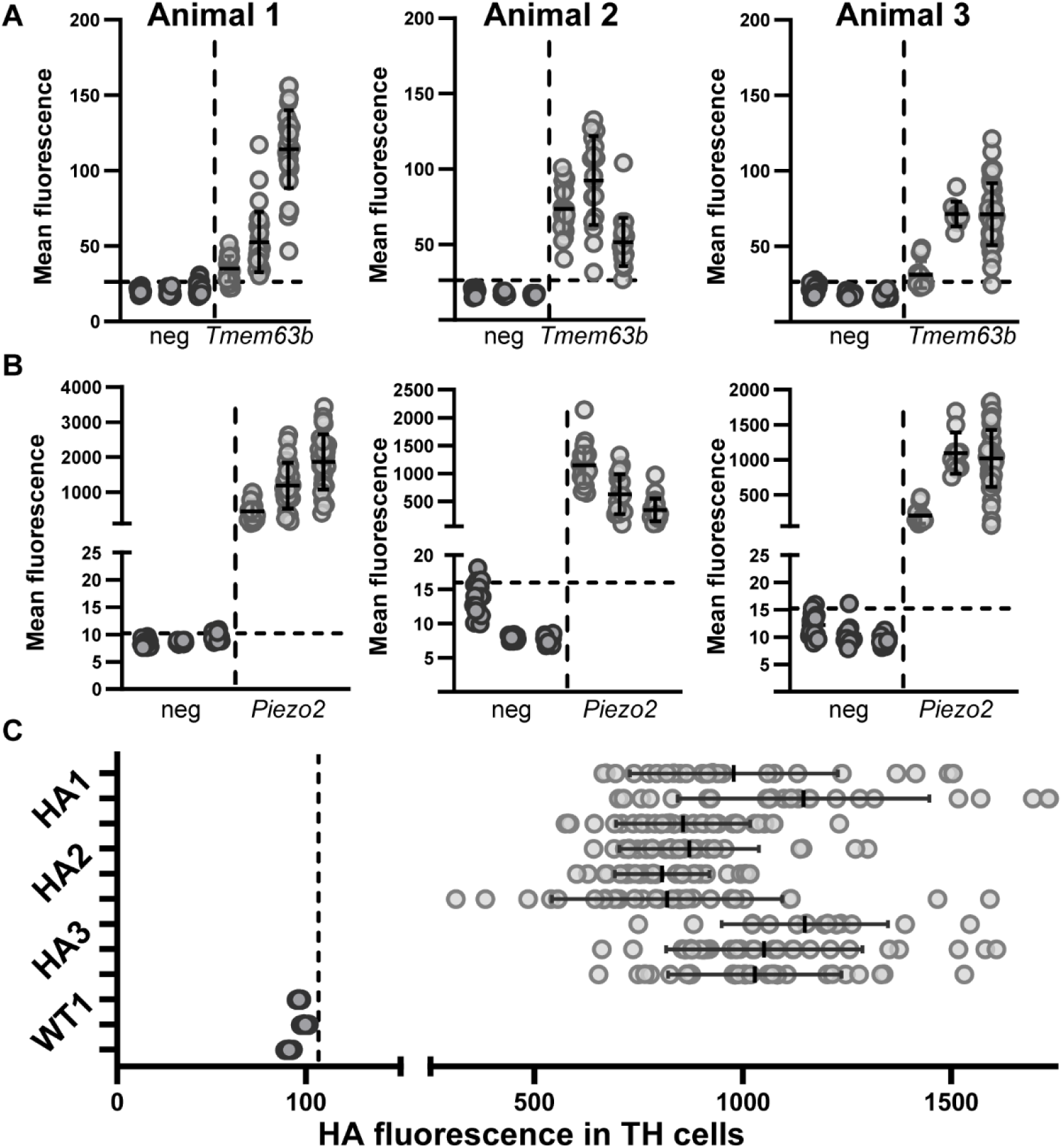
Negative controls used to create threshold for ISH and IHC experiments. **A**: Each tdTomato positive neuron’s mean fluorescence from the *in situ* probes against *Tmem63b* and negative control probes across each section of DRG imaged for all three animals in the cohort. **B**: Same as above but *in situ* probe is for *Piezo2*. **C**: Mean fluorescence of HA antibody staining of TH positive DRG neurons in *Tmem63b^HA/HA^* vs *Tmem63b^WT/WT^*. Threshold made from the 99^th^ percentile of mean fluorescence of HA staining from *Tmem63b^WT/WT^* DRG used as a cut off for *Tmem63b^HA/HA^* HA staining. Bars represent mean ± S.D. Each dot represents a cell marked with TH antibody, 3-HA replicate per animal, N = 3 animals. 3 WT replicates, N = 1 animal.

